# Thermodynamic Cycles of Evolution: Specific Dissipation, Ecological Recovery, and Evolutionary Memory

**DOI:** 10.64898/2026.11.23.768549

**Authors:** Eva Déli, Edina Molnár, Reza Rastmanesh

**Affiliations:** Department of Psychology, Faculty of Health Sciences, University of Debrecen, Debrecen 4032, Hungary; The Nutrition Society, Boyd Orr House, 10 Cambridge Court, 210 Shepherds Bush Road, London W6 7NJ, UK

**Keywords:** specific dissipation, entropy production, ecological recovery, evolutionary memory, dormancy, rate–yield trade-off, consumer–resource model, mass extinction, maximum power principle

## Abstract

In the post-extinction period, does evolutionary capacity recover together with biomass? We address this question with a consumer–resource model in which strategies lie on a rate–yield (opportunist–gleaner) trade-off and dissipate the substrate free energy they do not conserve in biomass. We track biomass-specific dissipation, *s*, a model-based proxy for specific entropy production. After a reset, abundant resources favor fast, low-yield strategies, and *s* is high; this rate selection resembles, but does not demonstrate, the maximum power principle. As resources are drawn down, competitive exclusion favors high-yield strategies and lowers *s*, provided that turnover is low and that affinity is lost faster than growth rate is gained along the trade-off. In finite populations, the high-dissipation phase returned after a reset only if a reservoir of costly strategies, here dormant propagules, had persisted since those strategies were last abundant; its return fell from 199 to 33 of 200 runs as the reservoir decayed. Biomass recovered in every run, and sooner when the high-dissipation phase failed to return. Ecological recovery and recovery of evolutionary capacity are therefore distinct. We state falsifiable predictions for calorimetric chemostat experiments with graded bottlenecks and controlled resting-stage survival.

## 1. Introduction

### 1.1. A Thermodynamic Tension in Evolution

Living systems are open, far-from-equilibrium structures that persist by degrading free energy [1,2]. Two optimization principles proposed for such systems appear to conflict. The maximum power principle (MPP) of Lotka [3] and Odum and Pinkerton [4] holds that selection favors systems that maximize the rate of useful energy transformation. Selection for minimal dissipation is often linked to Prigogine’s theorem that linear, near-equilibrium steady states minimize entropy production (MinEP) [5]. A third principle, maximum entropy production (MaxEP) [6,7], is frequently conflated with the MPP but makes different claims. Ecology supports both tendencies: rate–yield trade-offs in microbial metabolism [8] and the replacement of fast-growing pioneers by efficient competitors during succession [9].

### 1.2. The Gap

The history of life is punctuated by extinctions, bottlenecks, and recoveries [10,11], and ecology describes similar recurrent dynamics at shorter scales: ecosystem development [9], r- and K-selection [12–14], ecological opportunity after extinction or colonization [15,16], and the adaptive cycle of panarchy theory [17,18]. Resource-competition theory predicts which strategies persist at high and low resource levels [19]. These frameworks leave two gaps. They lack a common physical quantity that would make the phases of recovery measurable and comparable, from microbial cultures to marine faunas [20]. And they treat recovery mainly as the return of biomass, diversity, or ecosystem function; they rarely ask whether a community that regains its biomass also regains the strategies it lost.

### 2.3. Question and Contribution

An earlier proposal described evolution as a recurrent cycle in which environmental upheavals remove lineages that cannot adapt and open space for new groups of organisms, which in turn transform their environment [21]. Here we model one part of that cycle: the shift from high-throughput to efficient strategies within a recovery, expressed in terms of specific dissipation and restricted to resource-limited systems with a rate–yield trade-off. Change across cycles remains a hypothesis (Section 6). We ask whether ecological recovery after a reset implies recovery of evolutionary capacity, defined here as the capacity of a community to re-express energetically costly strategies. We answer it in a minimal consumer–resource model with explicit dissipation, analyzed deterministically and with finite populations. Biomass-specific dissipation serves as the marker: it is high when costly, fast-growing strategies dominate and low when efficient competitors do. We distinguish three levels of claim:

1. Results of established resource-competition theory: which strategies persist at high and low resource levels (Section 4.2);
2. Results of our model under its assumptions: the condition under which competitive exclusion minimizes specific dissipation, the lag of beneficial substitution after a reset, and the decoupling of biomass recovery from the return of the high-dissipation phase (Sections 4.2–4.6);
3. Hypotheses: interpretations of the fossil record and a cross-cycle ratchet in energetic capacity (Sections 5 and 6).

The framework applies to resource-limited systems with a rate–yield trade-off. It is not a general law of evolution, and thermodynamics constrains the outcome without directing it.

## 2. Thermodynamic Framework

### 2.1 Entropy Production and the Dissipation Proxy

The principles at issue concern the rate of entropy production, σ (W·K^−1^), not the entropy *S* of a state. The two are independent: a low-entropy state can produce entropy rapidly, and a high-entropy state can be nearly inert. For a community of strategies *i* with biomass *B*_*i*_, total entropy production is σ = Σ_i_ σ_i_, and biomass-specific entropy production is *s* = σ/Σ_i_*B*_*i*_ (W·K^−1^·kg^−1^). We take the system to be a well-mixed culture vessel or habitat at constant temperature *T* _env_ that exchanges substrate, products, and biomass with its surroundings. The quantity we model (Equation (3)) is the rate at which substrate free energy not conserved in biomass is dissipated as heat within this boundary, divided by *T* _env_. It is therefore one component of the entropy balance: entropy carried across the boundary by matter flows is not tracked. We call *s* specific dissipation and use it throughout as a model-based proxy for biomass-specific entropy production, not as a complete entropy balance.

In practice, *s* is estimated from specific heat dissipation (W·kg^−1^), measured by calorimetry, divided by environmental temperature. This estimate is valid when entropy exchanged with matter flows is small relative to heat flow, as in aerobic, enthalpy-driven growth. It fails for strongly entropy-driven metabolisms, where heat production is a poor proxy for entropy production [22]. Ecological occupancy, the fraction of resource supply captured by resident populations, is used as a descriptive variable; extinctions lower occupancy, not thermodynamic entropy.

### 2.2. Three Principles That Are Often Conflated

Table 1 separates the three principles. Literal reconciliation of the MPP with MinEP is ill-posed, because their domains of validity do not overlap. Ecological equilibrium (a stationary population size) is not thermodynamic equilibrium, and a saturated community still processes large energy fluxes. The defensible claim concerns specific dissipation: mature ecosystems tend toward higher biomass and total throughput but lower dissipation per unit biomass [9,23]. Throughout, “minimal dissipation” means selection for low *s* and high yield, an ecological outcome of competition and not an instance of Prigogine’s theorem.

**Table 1.** Three optimization principles and their domains of validity.

| **Principle** | **Quantity Optimized** | **Domain of Validity** | **Ecological Reading** |
|---|---|---|---|
| Maximum power (MPP) [3,4] | Rate of useful work or energy capture | Open systems competing for a flow; optimum efficiency below maximum | Selection for fast acquisition and growth when resources are abundant |
| Maximum entropy production (MaxEP) [6,7] | Total *σ*, subject to constraints | Non-equilibrium steady states with many degrees of freedom; status debated | Ecosystems as dissipative structures [1] |
| Minimum entropy production (MinEP) [5] | Total *σ* at a steady state | Linear regime near equilibrium only | Not literally applicable to ecosystems, which are far from equilibrium |

Maximum population growth rate and maximum power are not equivalent: power is a rate of energy transformation, and in the original formulation it is maximal at intermediate efficiency [4]. Our model contains no such optimum (Section 4.2). What it shows is resource-dependent rate–yield selection: fast, low-yield strategies win when resources are abundant, and high-yield strategies win when resources are limiting. We call the first regime *MPP-like selection* to mark its resemblance to the MPP, which serves here as theoretical background, not as a mechanism the model demonstrates.

### 2.3. The Cycle

The cycle has four phases, labeled with panarchy notation (Section 3):

1. Reorganization (α): after an extinction, colonization, or other reset, occupancy is low, and resource supply exceeds demand;
2. Exploitation (r): strategies with high maximal growth rate, high throughput, and high *s* spread under MPP-like selection;
3. Conservation (K): as resources are drawn down, competitive exclusion favors strategies that persist at the lowest resource level, typically high-yield, low-*s* specialists;
4. Release (Ω): a perturbation removes a large fraction of biomass and resets occupancy.

Transitions are governed by measurable control parameters: resource level relative to the competitors’ half-saturation constants, turnover, and the frequency and duration of perturbations. The cycle itself is ecological; evolutionary change (mutation, selection, and the loss or retention of strategies) takes place within it and is what the model tracks.

## 3. Relation to Existing Frameworks

### 3.1. The Adaptive Cycle

The adaptive cycle [17,18] describes systems by their potential (accumulated capital such as biomass), connectedness, and resilience as they pass through exploitation (r), conservation (K), release (Ω), and reorganization (α). Panarchy links nested cycles across scales; in its “remember” linkage, a larger, slower level supplies material for renewal, such as seed banks, refugia, and surviving lineages. Sundstrom and Allen [20] noted that quantitative measures of the cycle’s dimensions remain scarce. Table 2 maps the cycle onto the present framework and names a measurable correlate for each phase.

**Table 2.**
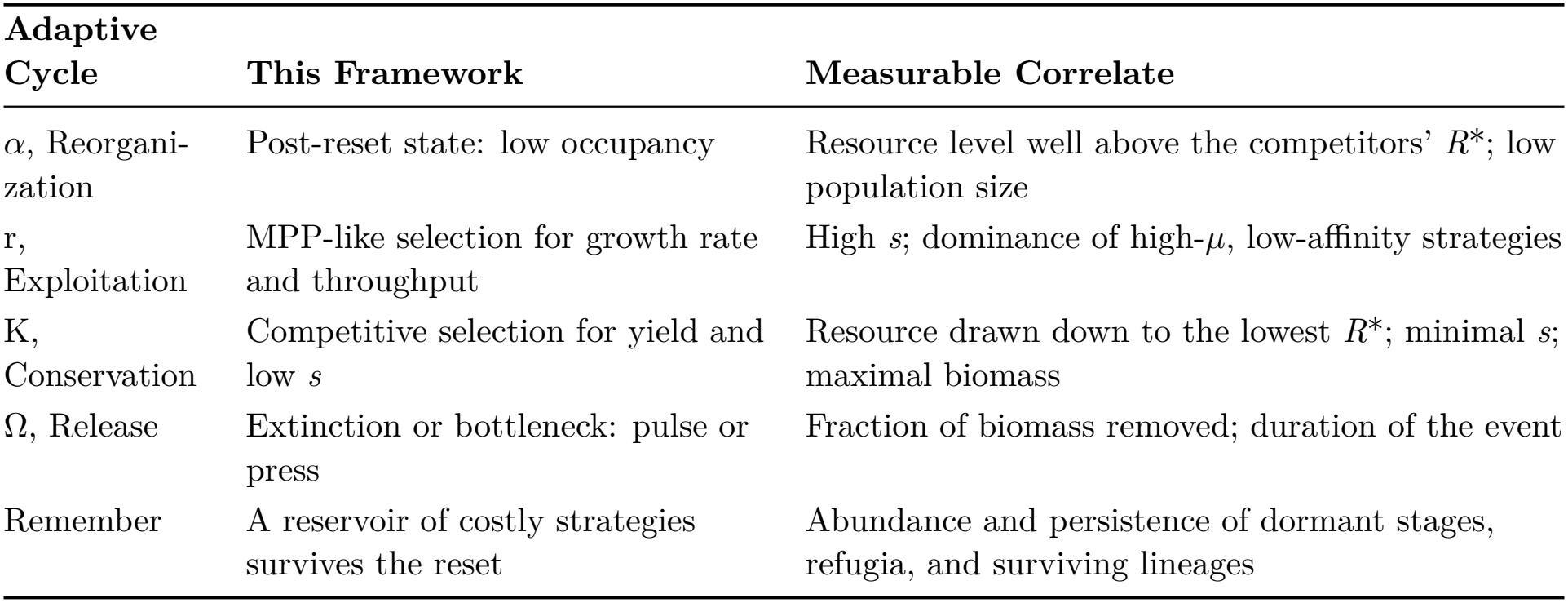
Correspondence between the adaptive cycle and the present framework. *R*\* is the break-even resource level (Equation (4)).

| **Adaptive Cycle** | **This Framework** | **Measurable Correlate** |
|---|---|---|
| α, Reorgani-zation | Post-reset state: low occupancy | Resource level well above the competitors' *R*<sup>\*</sup>; low population size |
| r, Exploitation | MPP-like selection for growth rate and throughput | High *s*; dominance of high-*μ*, low-affinity strategies |
| K, Conservation | Competitive selection for yield and low *s* | Resource drawn down to the lowest *R*<sup>\*</sup>; minimal *s*; maximal biomass |
| Ω, Release | Extinction or bottleneck: pulse or press | Fraction of biomass removed; duration of the event |
| Remember | A reservoir of costly strategies survives the reset | Abundance and persistence of dormant stages, refugia, and surviving lineages |

### 3.2. What Thermodynamics Adds, and What It Does Not

Several conclusions below follow from classical theory and require no thermodynamics: the strategy with the lowest break-even resource level excludes the others [19], and fast strategies precede efficient ones during succession [9,14]. The thermodynamic description adds three things.

First, a physical currency with a measurement route. Specific dissipation can be measured in a chemostat by calorimetry combined with yield, bounded for fossil faunas by body size and metabolic scaling [24,25], and compared across scales; panarchy’s potential and connectedness have no such unit.

Second, a quantity distinct from biomass-specific metabolic rate. Specific metabolic rate measures total energy turnover per unit biomass; specific dissipation excludes the free energy retained in biomass and is bounded below by the thermodynamic limit on yield (Equation (5)). The two are ordered alike at steady state, but *s* separates strategies more sharply: in the baseline parameterization, the steady-state contrast between opportunist and gleaner is about ninefold in *s* and under threefold in specific metabolic rate.

Third, conditions that the ecological description does not state: when competitive exclusion minimizes *s*, and the turnover rate above which it does the opposite (Section 4.2); and an energetic signature, the return or absence of a high-dissipation phase, of whether costly strategies survived a reset (Section 4.6).

If these predictions fail, the thermodynamic vocabulary is a relabeling of succession and should be dropped.

## 4. A Minimal Model

The model has an ecological layer with explicit dissipation and an extinction operator. We analyze it deterministically (Sections 4.1–4.5) and with finite populations (Section 4.6).

### 4.1. Ecological Layer with Explicit Dissipation

Consider *n* strategies competing for one limiting substrate of concentration *R* (mol·m^−3^) in an open system with supply concentration *R*_0_ and turnover rate *D* (s^−1^), as in a chemostat. Strategy *i* has biomass concentration *N*_*i*_ (kg·m^−3^), maximal growth rate µ_i_, half-saturation constant *K*_*i*_, and yield *Y*_*i*_ (kg·mol^−1^):

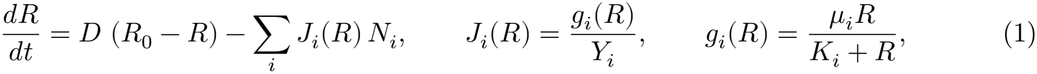

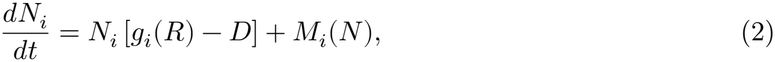

where *J*_*i*_ is the specific uptake rate (mol·kg^−1^·s^−1^), *g*_*i*_ is the specific growth rate, and *M*_*i*_ is nearest-neighbor mutation at rate *u*. Strategies lie on a gleaner–opportunist trade-off axis θ ∈ [0, 1] [26], for which the rate–yield trade-off of ATP-producing pathways provides a biochemical basis [8]. Opportunists (θ = 1) have high µ, low affinity (high *K*), and low yield; gleaners (θ = 0) have the opposite.

Let Δ*G* (J·mol^−1^) be the free energy released per mole of substrate consumed. We approximate the fraction conserved in biomass by *Y*_*i*_/*Y* _max_, where *Y* _max_ is the thermodynamic upper bound on yield; the remainder is dissipated:

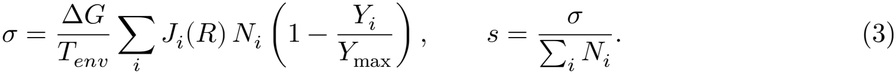

With these units, σ is in W·K^−1^·m^−3^ and *s* in W·K^−1^·kg^−1^. Equation (3) defines the dissipation proxy; it is phenomenological, not a derived balance. Yield is a mass ratio, and the conserved fraction of free energy also depends on biomass composition, substrate oxidation state, maintenance, and electron acceptor [22]. We therefore tested a nonlinear allocation, with conserved fraction (*Y*_*i*_/*Y* _max_)^*p*^, *p* = 0.5–3 (*p* = 1 is Equation (3)). The conclusions below require only that the dissipated fraction decreases with yield (Section 4.2), and they held across this range (Table S4).

### 4.2. Two Selective Regimes

#### MPP-like regime

Immediately after a reset, *R* ≈ *R*_0_ ≫ *K*_*i*_, so *g*_*i*_ ≈ µ_i_ and the fastest-growing strategy spreads. Per unit biomass at *R*_0_, throughput *J*_*i*_, conserved (useful) power Δ*G J*_*i*_*Y*_*i*_/*Y* _max_, and dissipation all rise monotonically from gleaner to opportunist (Figure 1b), so the axis has no power optimum at intermediate efficiency, unlike the original MPP [4]; the maximum lies at the end of the frontier. At steady state, specific useful power is the same for all strategies (Δ*G D*/*Y* _max_), and total useful power is highest for the high-yield strategy, which supports the largest biomass *B* = *Y* (*R*_0_ − *R*\*). The model thus demonstrates rate selection with high specific dissipation, not maximum power.

**Figure 1.**
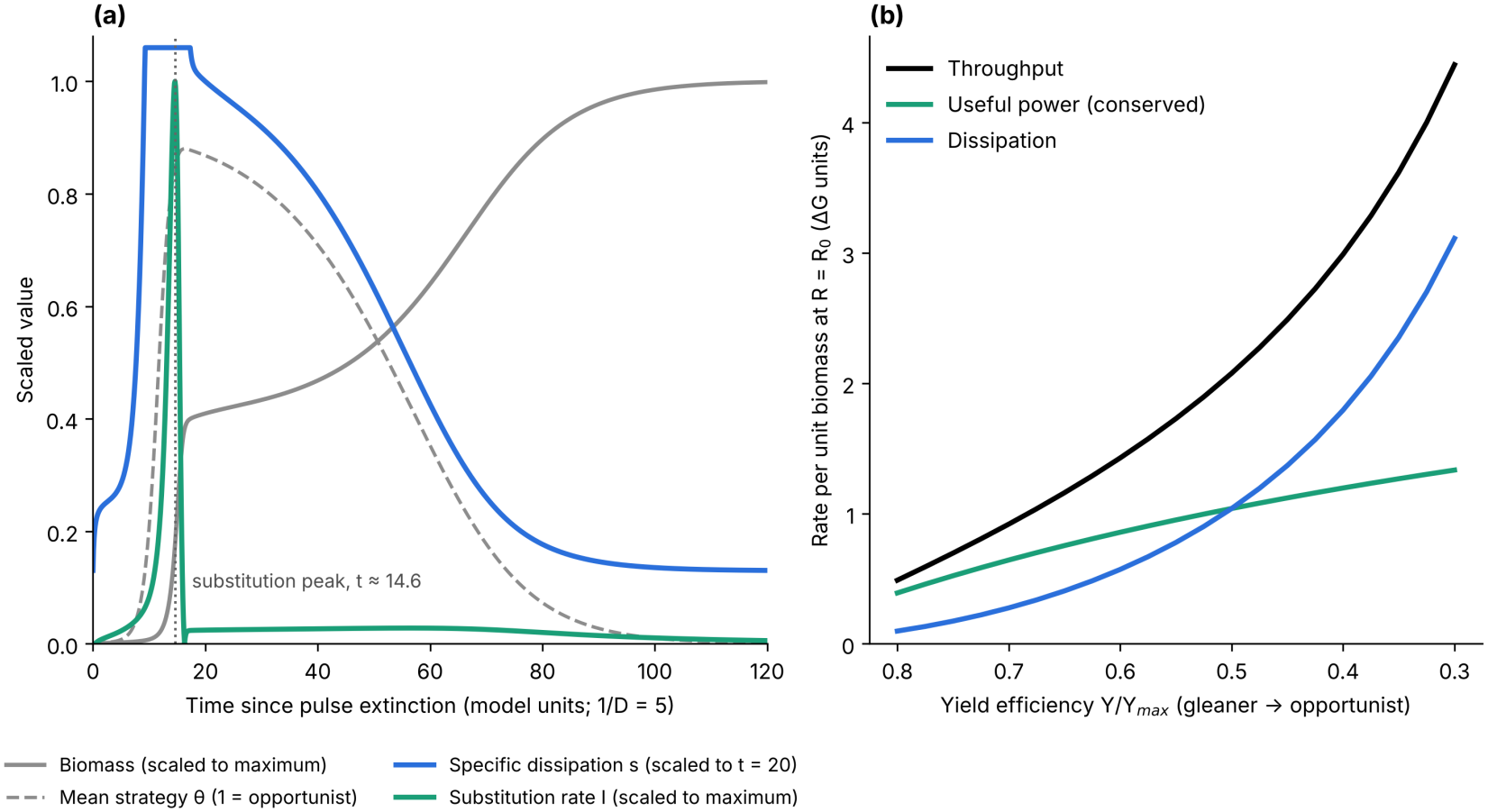
(**a**) Deterministic trajectory after a pulse extinction (*f* = 10^−3^) at *t* = 0. Biomass and beneficial substitution rate *I* are scaled to their maxima, *s* to its level at *t* = 20, and θ is the mean strategy (1 = opportunist). *s* exceeds the plotted range (to about 5 times its level at *t* = 20) during exponential growth and then falls about eightfold as gleaners replace opportunists.(**b**) Throughput, conserved (useful) power, and dissipation per unit biomass at *R* = *R*_0_ across the trade-off axis, in units of Δ*G*; none has an interior maximum.

#### Conservation regime

At steady state, strategy *i* persists only above its break-even resource level,

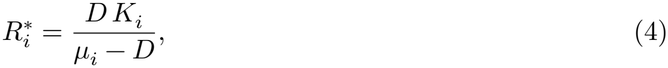

and the strategy with the lowest *R*\* excludes the others [19]. This result alone does not establish that the survivor minimizes dissipation. At the steady state of a resident *i, g*_*i*_(*R*_*i*_*) = *D*, and Equation (3) gives

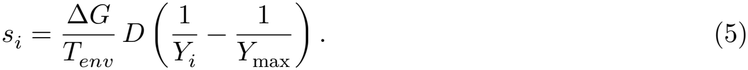

Steady-state specific dissipation therefore depends on the strategy only through its yield: *s*_*i*_ < *s*_*j*_ if and only if *Y*_*i*_ > *Y*_*j*_. The implication *R*_*i*_* < *R*_*j*_* ⟹ *s*_*i*_ < *s*_*j*_ holds exactly when break-even resource level and yield are inversely ordered across strategies. On a trade-off axis along which yield declines, this requires d[*K* /(µ − *D*)]/dθ > 0, that is, *K* ^′^/*K* > µ^′^/(µ − *D*): affinity must be lost faster than net growth rate is gained. For the linear trade-off used below, the condition holds for *D* < *D*_*c*_ = 0.316. Above *D*_*c*_ the ranking reverses: the opportunist has the lowest *R*\*, and competitive exclusion selects the strategy with the highest *s*. The value of *D*_*c*_ belongs to this parameterization; only the condition is general.

The size of the effect depends on the trade-off structure. With *K* ∝ θ^2^ the exclusion winner is intermediate (θ = 0.15) and *s* falls only 1.1-fold after a pulse, against about eightfold for the linear trade-off; with *K* ∝ θ^0.5^ the gleaner still wins (8.6-fold); 11 or 41 strategies gave results like those of 21 (Table S4). The direction of the fall in *s* is therefore robust wherever the condition holds, but its magnitude is not, and Prediction P1 (Section 7) concerns the direction.

In both regimes, substrate throughput *D*(*R*_0_ − *R*\*) approaches the supply limit *DR*_0_: selection changes what is done with the flux, not how much is captured. MPP-like selection and minimal specific dissipation are optima of different resource regimes, set by occupancy.

### 4.3. Extinction Events

We compare three events of equal depth *f*. A *pulse* multiplies every strategy’s active population by a survival fraction *f*, at random with respect to strategy. A *press* lowers the supply concentration so that active biomass falls to the fraction *f* of its conservation-phase value, holds it there for a duration *T*_*p*_, and then restores it; because *R*\* does not depend on *R*_0_, this press changes population size but not the selective ranking of strategies. A *delayed pulse* is a pulse applied after a waiting time *T*_*p*_; it controls for the time that elapses during a press.

### 4.4. Beneficial Substitution Rate and the Recovery Lag

We define the community beneficial substitution rate as the rate at which beneficial mutants are produced and fixed:

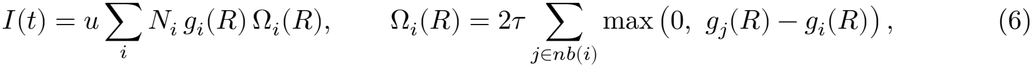

where the selective opportunity Ω_i_ sums the fixation probabilities of fitter neighboring strategies, each approximated by twice its selection coefficient *s*_sel_ = τ (*g*_*j*_ − *g*_*i*_). The reproductive timescaleτ (here 1 time unit) makes *s*_sel_ and Ω dimensionless. This is Haldane’s approximation, valid for weak selection in a population of roughly constant size (1/*N*_*e*_ ≪ *s*_sel_ ≪ 1); it underestimates fixation in growing populations. We use *I* only to locate the timing of its peak, which is set by the product of mutational supply and selective opportunity rather than by the fixation formula: using the relative advantage per generation, (*g*_*j*_ − *g*_*i*_)/*g*_*i*_, changed the simulated lags by at most 0.2 time units. *I* counts beneficial substitutions, not novel functions, new metabolic pathways, or phenotypic disparity. Mutational supply rises with *N* after a reset, whereas selective opportunity collapses when the resource is drawn down; their product has a maximum.

To locate it, suppose that one strategy with net growth rate *r* = *g*(*R*_0_) − *D* leads regrowth from the surviving fraction *f* of the conservation-phase population *N*_*K*_. While *R* ≈ *R*_0_, *N* (*t*) = *fN*_*K*_e^*rt*^ and Ω is constant, so *I* grows exponentially. Differentiating Equation (6) for this strategy gives d*I* /d*t* = *I* [*g*(*R*) − *D* + (∂ln(*g*Ω)/∂*R*)(d*R*/d*t*)]. The bracket equals *r* while *R* ≈ *R*_0_. Once drawdown begins, d*R*/d*t* < 0 and both *g* and Ω fall with *R*, so the bracket changes sign and *I* peaks shortly after draw-down begins; we verify this numerically below. Draw-down begins when uptake *JN* becomes comparable to supply *DR*_0_, that is, when *N* reaches a fraction k of *N*_*K*_ that does not depend on *f*. Hence

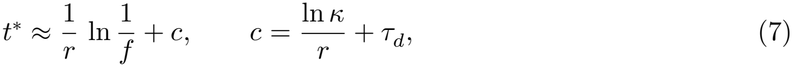

where τ _d_ is the duration of draw-down. The assumptions are *f* ≪ 1, a single strategy leading regrowth, and *R* ≈ *R*_0_ until *N* ≈ k*N*_*K*_. Equation (7) predicts the slope of the lag against ln(1/*f*), 1/*r*, and not only its logarithmic form.

### 4.5. Numerical Illustration

We simulated 21 strategies with µ from 0.4 to 2.0, *K* from 0.5 to 10, and *Y* from 0.8 to 0.3, all linear in θ; *Y* _max_ = 1; *D* = 0.2; *R*_0_ = 20; *u* = 10^−4^ (arbitrary units; Supplementary File S1). In this deterministic version, a minimum density of 10^−9^ per strategy acts as an implicit, inexhaustible reservoir, which Section 4.6 replaces with an explicit, finite one.

After a pulse (*f* = 10^−3^) imposed at steady state (Figure 1), opportunists dominated within about 20 time units (mean θ = 0.87). Specific dissipation spiked transiently to about five times its later plateau during exponential growth, settled near 0.39, and then fell about eightfold to 0.05 over the next ∼80 time units as gleaners replaced opportunists, in agreement with Equation (5). Biomass rose from 6.4 to 15.6. Total throughput stayed near the supply limit (*DR*_0_ = 4), while total dissipation fell from 2.5 to 0.8 and conserved power rose from 1.3 to 3.1 (Δ*G* units), because higher yield diverts more of a fixed flux into biomass. In open food webs, total dissipation can rise during succession [1,9] while specific dissipation still falls. The robust prediction is therefore the trajectory of *s*, not that of σ.

The beneficial substitution rate peaked 14.6 time units after the pulse. Across severities from *f* = 10^−1^ to 10^−7^, the lag rose from 9.5 to 21.7 time units, linearly in ln(1/*f*) with a slope of 2.1 time units per tenfold increase in severity (2.3 over 10^−1^–10^−5^). Equation (7) predicts 2.0 for regrowth led by the fastest strategy (*r* = 1.13); the difference reflects the simultaneous regrowth of intermediate strategies. The peak followed the onset of net draw-down (the maximum of *R*) by 3.0–4.1 time units at every severity, as the derivation requires.

### 4.6. Finite Populations, Dormancy, and Recovery of Function

To test which conclusions survive demographic noise, we implemented a stochastic version of Equations (1) and (2) (Supplementary File S2). Individuals are counted explicitly (about 15,000 in the conservation phase); births (Poisson), deaths and dormancy (binomial), and mutations are random events advanced by tau-leaping with a time step of 0.02; and there is no density floor (NumPy PCG64 generator; seeds are in the code). A lost strategy returns only by mutation or from an explicit bank of dormant propagules. Active individuals enter dormancy at rate δ = 0.01, and propagules reactivate at rate *a* = 0.002 and are lost at rate λ = 0.005, so that their persistence time is 1/(λ + *a*) ≈ 143. Each run starts from colonization and reaches the conservation phase before the event. We ran 200 replicates per condition and report proportions with 95% Wilson confidence intervals (CI). A high-dissipation phase is scored when peak *s* exceeds five times its conservation-phase value; gleaners regrowing alone reach about twice that value.

#### Active variation alone carries no memory

Without a bank, none of 200 runs showed a high-dissipation phase after a pulse (*f* = 10^−2^). Opportunists lie 10–20 mutational steps from the gleaner, and the window of abundant resource closes before mutation reaches them. Biomass is nevertheless recovered (median 25.5 time units). Without a bank in gleaner communities, the median lag of the substitution peak rose from 4.0 to 35.5 time units for *f* from 0.3 to 10^−3^, a slope of 12.6 per tenfold severity against 12.1 predicted by Equation (7) for gleaner-led regrowth (*r* = 0.19), independent of the density floor of Section 4.5.

#### A reservoir restores the high-dissipation phase

With a bank, a pulse that spared dormant stages was followed by a high-dissipation phase in 199 of 200 runs (CI 0.97–1.00), with peak *s* at a median of 18.5 times its conservation-phase value (Table 3, Figure 2a). When the pulse killed dormant stages at the same rate as active ones, the phase returned in 58 of 200 runs (CI 0.23–0.36). In this model, the MPP-like phase therefore requires a reservoir of costly strategies, which makes the “remember” linkage of panarchy an explicit model component.

**Table 3.** Events of matched depth (*f* = 10^−2^) in the finite-population model with propagule loss rate λ = 0.005; 200 runs per row. A high-dissipation phase is a peak *s* above five times the conservation-phase value. Times are in model units after the event ends. CI: Wilson confidence interval; IQR: interquartile range.

| **Event** | **Waiting Time *T<sub>p</sub>*** | **Opportunist Propagules at Reset** | **Runs with High-Dissipation Phase (95% CI)** | **Peak *s* Relative to Conservation Phase, Median (IQR)** | **Biomass Recovery Time, Median (IQR)** |
|---|---|---|---|---|---|
| Pulse | 0 | 792 | 199/200 (0.97–1.00) | 18.5 (15.5–21.0) | 51.0 (48.9–53.0) |
| Press | 150 | 278 | 200/200 (0.98–1.00) | 17.2 (14.6–19.3) | 51.5 (49.0–53.0) |
| Delayed pulse | 150 | 286 | 198/200 (0.96–1.00) | 15.5 (12.3–18.2) | 46.5 (44.9–49.0) |
| Press | 300 | 101 | 183/200 (0.87–0.95) | 16.1 (10.4–22.1) | 55.5 (50.5–59.0) |
| Delayed pulse | 300 | 102 | 134/200 (0.60–0.73) | 9.4 (3.6–14.5) | 40.0 (25.8–46.0) |
| Press | 600 | 13 | 70/200 (0.29–0.42) | 2.0 (1.9–10.3) | 29.0 (26.9–52.6) |
| Delayed pulse | 600 | 13 | 33/200 (0.12–0.22) | 1.9 (1.9–2.3) | 21.5 (21.5–22.5) |
| Pulse, no bank | 0 | – | 0/200 (0.00–0.02) | 1.9 (1.9–1.9) | 25.5 (25.0–26.0) |

**Figure 2.**
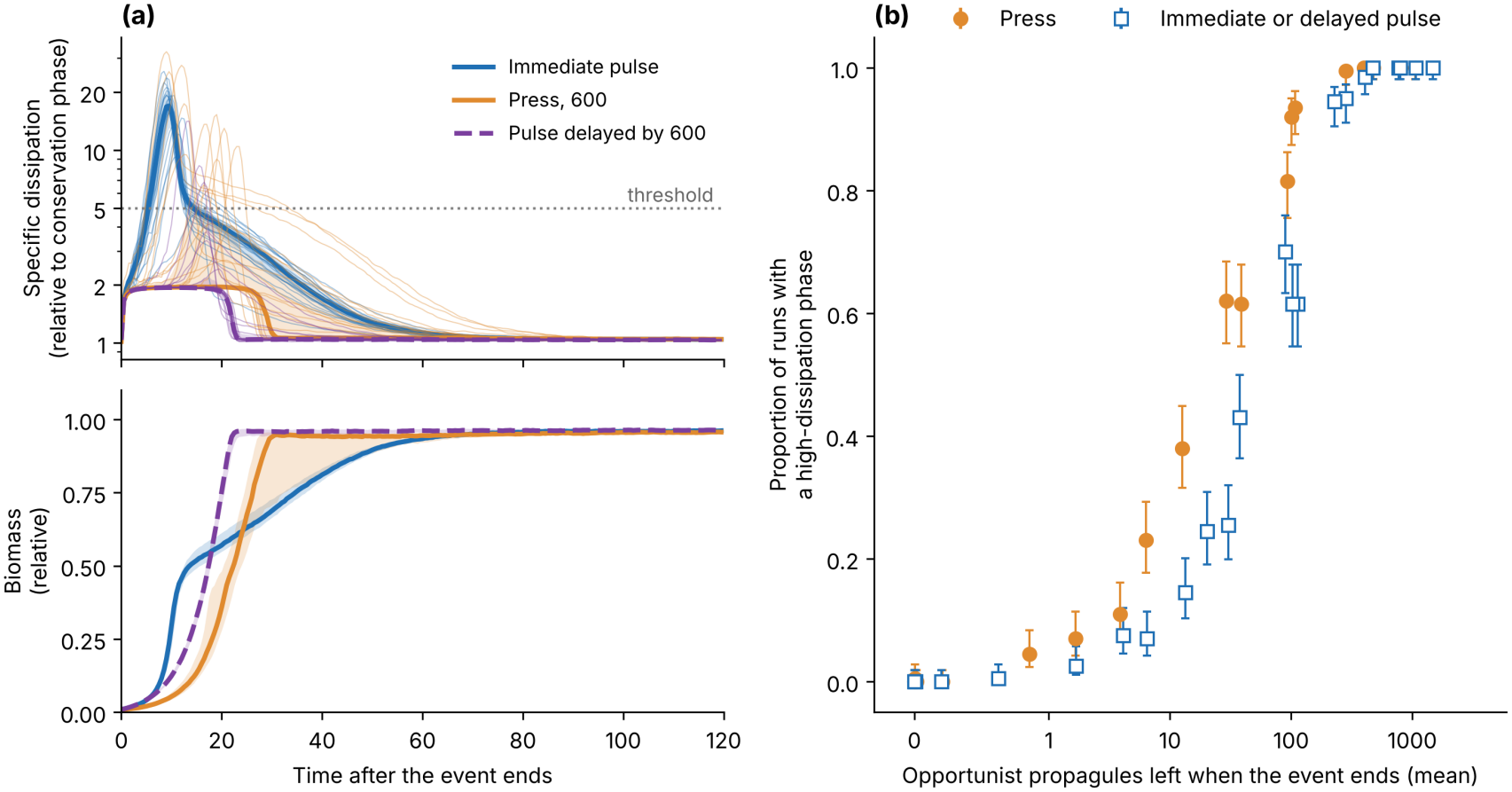
Recovery in the finite-population model after events of matched depth (*f* = 10^−2^). (**a**) Specific dissipation relative to its conservation-phase value (top; logarithmic scale) and biomass relative to its conservation-phase value (bottom) after an immediate pulse, a press of 600 time units, and a pulse delayed by 600 time units (λ = 0.005); thick lines are medians, bands are interquartile ranges of 200 runs, and thin lines are 20 individual runs. Biomass recovers in all three cases; the high-dissipation phase does not. (**b**) Proportion of runs with a high-dissipation phase against the mean number of opportunist propagules (θ ≥ 0.5) remaining when the event ends, for presses and for immediate or delayed pulses across four propagule loss rates (λ = 0.0025–0.02) and waiting times up to 900 time units; bars are 95% Wilson confidence intervals.

#### Memory decays with the time since costly strategies were last abundant

When the reset came later, at the end of a press of duration *T*_*p*_ or as a pulse delayed by *T*_*p*_, the opportunist reservoir had decayed, from 792 propagules to 13 after 600 time units, and the high-dissipation phase became rarer (Table 3). Across four loss rates (λ = 0.0025–0.02) and waiting times up to 900 time units, the outcome was governed by the number of opportunist propagules remaining at the reset, regardless of which combination of loss rate and waiting time produced it (Figure 2b). With faster loss (λ = 0.02), the phase followed only 56 of 200 immediate pulses (CI 0.22–0.35; Table S1).

#### Under the modeled press, elapsed time matters more than event type

With equal waiting time and equal reservoirs, the phase returned more often after a press than after a delayed pulse: 183 versus 134 of 200 runs at *T*_*p*_ = 300 (Fisher’s exact test, *p* < 0.001; risk difference 0.24, 95% CI 0.17–0.32) and 70 versus 33 at *T*_*p*_ = 600 (*p* < 0.001; risk difference 0.18, 95% CI 0.10–0.27). A press leaves alive the few opportunists that have just reactivated from the bank (a mean of 1.7 individuals at *T*_*p*_ = 300), whereas a pulse kills 99% of them. This asymmetry holds for a press that lowers supply without changing the selective ranking; it does not show that press disturbances are generally less damaging than pulses (Section 5.3).

#### Biomass recovery is not recovery of function

Biomass regained 90% of its conservation-phase level in every run of the conditions in Tables 3 and S1. It did so sooner when the high-dissipation phase failed to return (median 21.5 time units after a pulse delayed by 600, against 51.0 after an immediate pulse), because gleaners regrew without a wasteful opportunist bloom (Figure 2a). A community can therefore regain its biomass while losing its capacity to express energetically demanding strategies.

#### What is built in, and what is not

That a reservoir preserves costly strategies is partly built into the model. The non-trivial results are the dependence on elapsed time, the press–pulse asymmetry, and the decoupling of biomass from function. We distinguish evolutionary memory in general, any means by which a community retains access to strategies it is not currently expressing, from the mechanism modeled here, persistence of dormant propagules. Refugia, migration, cryptic variation, recombination, and horizontal transfer could serve similarly and were not simulated, although active variation alone proved insufficient. The ordering of the three events and the faster biomass recovery without the phase held in all 12 variants of a one-at-a-time sensitivity analysis (each dormancy, population-scale, mutation, or pulse-depth parameter changed two- to tenfold up and down; Table S2), under alternative thresholds (Table S3), and under the allocation-independent criterion that mean θ exceeds 0.5 (157 of 200 immediate pulses; 0 of 200 without a bank). Nonlinear allocation (*p* = 0.5–3) leaves the dynamics unchanged and alters only how *s* is scored (Table S4). The phase also requires a deep reset: after a milder pulse (*f* = 0.1) it followed only 38 of 200 pulses (CI 0.14–0.25). Halving the time step and rerunning with two independent sets of random seeds reproduced all qualitative results; the press–delayed-pulse risk difference ranged from 0.16 to 0.26 across waiting times and reruns.

### 4.7. Implementation

The deterministic and finite-population models were implemented in Python (Files S1 and S2). The code was written and run with the assistance of an AI model (Sonnet 5.5). The authors specified the model equations, parameters, and analyses. One author (E.D.) independently reran all analyses except the regime diagram of Figure 2b on a separate computer and reproduced the deposited outputs exactly. Robustness to the numerical time step and to the random seeds was also checked.

## 5.3 Comparison with Evidence

The comparisons in this section are illustrative and hypothesis-generating: none of the data measures dissipation, and each pattern has alternative explanations.

### 5.1. Delayed Recovery in the Fossil Record

Kirchner and Weil [27] found that origination peaks lag extinction peaks by roughly 10 Myr throughout the Phanerozoic, with comparable lags after large and small extinctions. The model predicts a lag that depends only logarithmically on severity (Equation (7)), qualitatively consistent with similar delays after large and small events; we claim no quantitative fit. With an assumed net diversification rate of 0.2 per lineage per Myr and 10% survival, the model lag is of order 10 Myr, an order-of-magnitude correspondence only.

### 5.2. The Rarity of Early Bursts

Harmon et al. [28] found little support for early-burst models of body size and shape evolution across many animal clades. Early-burst tests on living species have low power [29], whereas fossil disparity often peaks early [30]. The framework predicts early bursts only in clades that diversified into low-occupancy ecospace; most clades in a broad sample diversified into occupied communities, where dynamics are expected to be constrained.

### 5.3. Pulse versus Press Extinction in Digital Communities

Yedid et al. [31] evolved communities of Avida digital organisms [32] under instantaneous pulse extinctions with random survivors and under press extinctions, prolonged periods of low resource availability. Functional activity and phenotypic diversity recovered faster after pulses, and adaptation during the press, toward short generation times and low functional activity, degraded the ability to re-evolve earlier functions [33]. Our model does not reproduce a press-specific delay, because its press leaves the selective ranking unchanged and its memory resides in a dormant bank; in Avida, the press changed the direction of selection and memory resided only in the active population. A press may therefore delay functional recovery by adaptation to the press environment or by outlasting the memory of costly strategies, not through scarcity itself.

### 5.4. Two Historical Illustrations

#### The Ediacaran–Cambrian radiation

The end-Ediacaran turnover (∼550–539 Ma) removed much of the Ediacaran biota [34,35], and benthic ecospace was then filled by mobile, burrowing, and predatory animals [36]. Rates of phenotypic and molecular evolution among Cambrian arthropods were about four to five times higher than in later intervals [37], disparity peaked early in Cambrian clades [30], and the high-turnover Cambrian fauna was succeeded by lower-turnover Paleozoic faunas dominated by suspension feeders with low metabolic rates [25,38,39]. This sequence is consistent with a shift from high to low specific dissipation, but no estimate of dissipation exists for these faunas, and oxygenation offers an alternative or complementary explanation [40–42].

#### Darwin’s finches

On Daphne Major, the 1977 drought reduced the medium ground finch (*Geospiza fortis*) by roughly 85% and favored larger, deeper beaks [43]; abundant small seeds after the 1982– 1983 El Niño reversed selection [44]; and competition with *G. magnirostris* during the 2003–2005 drought caused character displacement toward smaller beaks [45]. These crashes and booms match the ecological phases of the cycle, but their thermodynamic interpretation is untested; it would require field metabolic rates (doubly labeled water) combined with seed-handling efficiency.

## 6. Limitations and Future Directions

The model considers one limiting substrate, a prescribed trade-off frontier, and a fixed environment, without trophic interactions or spatial structure. The cycle should therefore be read as a conditional framework for resource-limited systems with a rate–yield trade-off. Natural extensions include deriving dissipation from substrate and biomass Gibbs energies, jointly exploring parameter space, and alternative memory mechanisms such as spatial refugia.

A cross-cycle hypothesis follows from the model but is not a result of it. Within a cycle, competition lowers specific dissipation, yet across the history of life the range of energetic designs has expanded in steps [46,47]. Release phases may be windows in which energetically costly designs can establish, with the lineages carrying them persisting through later collapses. Because the model holds the trade-off frontier fixed, it cannot generate new designs. The hypothesis predicts that first appearances of energetically demanding innovations, such as endothermy [48,49], are more frequent in recovery intervals than in matched background intervals; Vermeij’s escalation hypothesis [50], in which such innovations arise in saturated ecosystems, predicts the opposite.

## 7. Testable Predictions

Table 4 lists four predictions with a test system and the result that would count against the frame-work. P1–P3 can be tested within a few years at modest cost in microbial or digital communities. Failure of P1 at low turnover would refute the central claim about specific dissipation; failure of P3 would refute the decoupling of ecological and evolutionary recovery; failure of P4 alone would refute only the extension to macroevolution.

**Table 4.** Testable predictions and falsification criteria.

|  | Prediction | Test System | Falsified If |
|---|---|---|---|
| P1 | Specific heat dissipation is highest in the exploitation phase and declines as resources are drawn down, while substrate throughput stays near the supply limit; above a critical turnover rate the ordering reverses (Section 4.2) | Chemostat or serial-transfer communities after dilution shocks; isothermal microcalorimetry with yield measurement | *s* does not decline at low turnover, or declines without a shift toward higher-yield, higher-affinity strategies |
| P2 | The beneficial substitution rate peaks after a lag that increases with ln(1/*f*) with slope 1/*r* (Equation (7)) | Microbial evolution with graded bottlenecks; digital evolution; time-resolved sequencing | No lag, or a lag that does not scale with the logarithm of bottleneck depth |
| P3 | After a reset, the high-dissipation phase returns only if the time since costly strategies were last abundant is short relative to the persistence of their reservoir; biomass recovers in either case | Microbial communities with measured resting-stage survival, reset by dilution or starvation at controlled intervals, with calorimetry; digital communities with and without dormancy | Return of the high-dissipation phase is independent of elapsed time and reservoir persistence, or biomass fails to recover whenever that phase fails to return |
| P4 | Early-burst signals are enriched in clades originating soon after release events or colonization of vacant habitats | Phylogenetic comparative data with fossil tips; crown ages relative to extinction boundaries | No enrichment after controlling for clade age and size |

## 8. Conclusions

Ecological recovery and recovery of evolutionary capacity are distinct. In a consumer–resource model with a rate–yield trade-off and explicit dissipation, biomass returned after every reset we simulated, but the high-dissipation phase, the signature of costly, fast-growing strategies, returned only when a reservoir of those strategies had persisted since they were last abundant. Biomass recovered sooner when that phase failed to return. A community can therefore regain its biomass while losing its capacity to express energetically demanding strategies.

Biomass-specific dissipation makes this distinction measurable. It is high under MPP-like rate selection when resources are abundant and falls as competitive exclusion favors high-yield strategies, provided that turnover is low and that affinity is lost faster than growth rate is gained. It separates strategies more sharply than specific metabolic rate does, and calorimetry with yield measurement can track it through a recovery.

The framework applies to resource-limited systems with a rate–yield trade-off and is not a general law of evolution. Its predictions can be tested in chemostat and digital communities with graded bottlenecks and controlled resting-stage survival. Thermodynamics constrains which strategies ecological conditions make selectable; it does not direct evolution toward an endpoint.

## Supporting information

https://doi.org/10.5281/zenodo.24346681

## Supplementary Materials

The following supporting information is provided with the submission: File S1: deterministic model (Python); File S2: finite-population model (Python); File S3: Table S1 (matched-depth events with fast propagule loss), Table S2 (sensitivity analysis), Table S3 (robustness to the outcome threshold), and Table S4 (robustness to the yield–dissipation allocation, trade-off curvature, and number of strategies).

## Author Contributions

Conceptualization, E.D.; methodology, software, formal analysis, writing—original draft preparation, E.D., E.M., and R.R.; writing—review and editing, E.D., E.M., and R.R. All authors have read and agreed to the published version of the manuscript.

## Data Availability Statement

No empirical data were generated. The simulation code (Files S1 and S2), parameter values, random seeds, generated outputs, and figure source data that reproduce all numerical results are openly available in Zenodo at https://doi.org/10.5281/zenodo.24346681.

## Acknowledgments

During the preparation of this manuscript, the authors used Sonnet 5.5 to draft and edit the text, write the simulation code, and run the simulations (Section 4.7). The authors have reviewed and edited the output and take full responsibility for the content of this publication.

## Abbreviations

The following abbreviations are used in this manuscript:

MPP: maximum power principle
MaxEP: maximum entropy production
MinEP: minimum entropy production
CI: confidence interval
IQR: interquartile range.

