## Supplementary material for "Thermodynamic Cycles of Evolution: Specific Dissipation, Ecological Recovery, and Evolutionary Memory": https://doi.org/10.5281/zenodo.24346681

### Supplementary File S3

Thermodynamic Cycles of Evolution: Specific Dissipation, Ecological Recovery, and Evolutionary Memory

All tables report the finite-population model of Section 4.6 (Supplementary File S2), with 200 runs per condition. A high-dissipation phase is a peak specific dissipation above five times the conservation-phase value, unless stated otherwise. CI: 95% Wilson confidence interval; IQR: interquartile range.

**Table S1.** Events of matched depth ($f = 10^{-2}$) with fast propagule loss ($\lambda = 0.02$). Columns as in Table 3 of the main text.

| Event | Waiting Time $T_p$ | Opportunist Propagules at Reset | Runs with High-Dissipation Phase (95% CI) | Peak $s$ Relative to Conservation Phase, Median (IQR) | Biomass Recovery Time, Median (IQR) |
|---|---|---|---|---|---|
| Pulse | 0 | 20 | 56/200 (0.22–0.35) | 2.0 (1.9–5.6) | 25.0 (24.5–41.0) |
| Press | 150 | 1 | 4/200 (0.01–0.05) | 1.9 (1.9–1.9) | 27.5 (26.0–28.1) |
| Delayed pulse | 150 | 1 | 0/200 (0.00–0.02) | 1.9 (1.9–1.9) | 25.0 (24.5–25.5) |
| Press | 300 | 0 | 1/200 (0.00–0.03) | 1.9 (1.9–1.9) | 28.5 (27.5–30.5) |
| Delayed pulse | 300 | 0 | 0/200 (0.00–0.02) | 1.9 (1.9–1.9) | 25.0 (24.0–25.5) |
| Press | 600 | 0 | 0/200 (0.00–0.02) | 1.9 (1.9–1.9) | 29.0 (27.5–30.5) |
| Delayed pulse | 600 | 0 | 0/200 (0.00–0.02) | 1.9 (1.9–1.9) | 25.0 (24.5–25.5) |

**Table S2.** One-at-a-time sensitivity analysis. Baseline: $\delta = 0.01$, $a = 0.002$, $\lambda = 0.005$, $\omega = 1000$ individuals per unit biomass, $u = 10^{-4}$, $f = 10^{-2}$. Entries are runs with a high-dissipation phase out of 200 (95% CI) after an immediate pulse, a press of 300 time units, and a pulse delayed by 300 time units, followed by the median biomass recovery time for the three events. The last column gives the two-sided Fisher's exact test of press against delayed pulse.

| Parameter Changed | Immediate Pulse | Press, 300 | Delayed Pulse, 300 | Biomass Recovery Time (Pulse/Press/Delayed) | Press vs. Delayed Pulse, $p$ |
|---|---|---|---|---|---|
| Baseline | 200/200 (0.98–1.00) | 182/200 (0.86–0.94) | 144/200 (0.65–0.78) | 51.0/55.0/40.5 | <0.001 |
| $\delta = 0.003$ | 192/200 (0.92–0.98) | 113/200 (0.50–0.63) | 73/200 (0.30–0.43) | 49.5/39.0/24.5 | <0.001 |
| $\delta = 0.03$ | 200/200 (0.98–1.00) | 200/200 (0.98–1.00) | 192/200 (0.92–0.98) | 92.5/115.0/71.5 | 0.007 |
| $a = 0.001$ | 200/200 (0.98–1.00) | 176/200 (0.83–0.92) | 138/200 (0.62–0.75) | 56.0/57.5/46.0 | <0.001 |
| $a = 0.004$ | 200/200 (0.98–1.00) | 168/200 (0.78–0.88) | 92/200 (0.39–0.53) | 45.0/51.5/27.8 | <0.001 |
| $\lambda = 0.0025$ | 200/200 (0.98–1.00) | 200/200 (0.98–1.00) | 191/200 (0.92–0.98) | 49.5/53.5/41.2 | 0.004 |
| $\lambda = 0.01$ | 190/200 (0.91–0.97) | 45/200 (0.17–0.29) | 15/200 (0.05–0.12) | 51.5/28.0/23.5 | <0.001 |
| $\omega = 300$ | 193/200 (0.93–0.98) | 104/200 (0.45–0.59) | 51/200 (0.20–0.32) | 47.5/42.2/22.2 | <0.001 |
| $\omega = 3000$ | 200/200 (0.98–1.00) | 200/200 (0.98–1.00) | 195/200 (0.94–0.99) | 52.5/58.5/45.5 | 0.061 |
| $u = 10^{-5}$ | 200/200 (0.98–1.00) | 180/200 (0.85–0.93) | 139/200 (0.63–0.75) | 51.0/55.0/40.8 | <0.001 |
| $u = 10^{-3}$ | 200/200 (0.98–1.00) | 189/200 (0.90–0.97) | 139/200 (0.63–0.75) | 51.0/55.8/40.5 | <0.001 |
| $f = 10^{-1}$ | 38/200 (0.14–0.25) | 24/200 (0.08–0.17) | 2/200 (0.00–0.04) | 23.0/21.8/14.5 | <0.001 |
| $f = 10^{-3}$ | 200/200 (0.98–1.00) | 194/200 (0.94–0.99) | 153/200 (0.70–0.82) | 58.0/62.5/46.0 | <0.001 |

**Table S3.** Robustness of Table 3 ($\lambda = 0.005$) to the criterion for a high-dissipation phase. Entries are numbers of runs out of 200 that meet each criterion: peak $s$ above 3, 5 (main text), or 10 times the conservation-phase value, and peak mean strategy $\theta$ above 0.5.

| Event | Waiting Time $T_p$ | Peak $s > 3\times$ | Peak $s > 5\times$ | Peak $s > 10\times$ | Peak Mean $\theta > 0.5$ |
|---|---|---|---|---|---|
| Pulse | 0 | 200 | 199 | 196 | 157 |
| Press | 150 | 200 | 200 | 192 | 147 |
| Delayed pulse | 150 | 200 | 198 | 172 | 98 |
| Press | 300 | 192 | 183 | 153 | 120 |
| Delayed pulse | 300 | 158 | 134 | 93 | 38 |
| Press | 600 | 82 | 70 | 53 | 36 |
| Delayed pulse | 600 | 41 | 33 | 16 | 6 |

**Table S4.** Robustness to the yield–dissipation allocation, the trade-off curvature, and the number of strategies. (a) Deterministic model (Section 4.5; pulse $f = 10^{-3}$, $D = 0.2$): conserved fraction of substrate free energy $(Y/Y_{\max})^p$; $K = 0.5 + 9.5\theta^q$; $M$ strategies. Spike: peak $s$ relative to its level 20 time units after the pulse; fall: that level relative to $s$ at the end of the run; winner: strategy $\theta$ with the lowest $R^*$ at $D = 0.2$; share of $D$: fraction of turnover rates in 0.005–0.395 for which $R^*$ increases along the whole axis. (b) Finite-population model (Section 4.6): runs out of 200 with a high-dissipation phase (peak $s > 5\times$ the conservation-phase value) for exponent $p$; the dynamics are identical across $p$, which changes only how $s$ is scored.

| Variant | Spike | Fall | Winner $\theta$ | Share of $D$ with Monotone $R^*$ |
|---|---|---|---|---|
| $p = 0.5$ | 5.2× | 9.3× | 0.00 | 80% |
| $p = 1$ (baseline) | 5.2× | 7.7× | 0.00 | 80% |
| $p = 2$ | 5.2× | 5.7× | 0.00 | 80% |
| $p = 3$ | 5.2× | 4.6× | 0.00 | 80% |
| $q = 0.5$ | 5.3× | 8.6× | 0.00 | 0% |
| $q = 2$ | 3.4× | 1.1× | 0.15 | 0% |
| $M = 11$ | 5.3× | 8.1× | 0.00 | 80% |
| $M = 41$ | 5.2× | 7.5× | 0.00 | 80% |

| Exponent $p$ | Immediate Pulse | Press, 300 | Delayed Pulse, 300 | Peak Mean $\theta > 0.5$ (Pulse/Press/Delayed) |
|---|---|---|---|---|
| 0.5 | 200/200 (0.98–1.00) | 183/200 (0.87–0.95) | 147/200 (0.67–0.79) | 162/115/36 |
| 1 | 200/200 (0.98–1.00) | 182/200 (0.86–0.94) | 144/200 (0.65–0.78) | 162/115/36 |
| 2 | 200/200 (0.98–1.00) | 178/200 (0.84–0.93) | 136/200 (0.61–0.74) | 162/115/36 |
| 3 | 199/200 (0.97–1.00) | 175/200 (0.82–0.91) | 130/200 (0.58–0.71) | 162/115/36 |
